# Convergent molecular evolution associated with repeated transitions to gregarious larval behaviour in Heliconiini

**DOI:** 10.1101/2025.01.14.633010

**Authors:** Francesco Cicconardi, Callum F. McLellan, Alice Seguret, W. Owen McMillan, Stephen H. Montgomery

## Abstract

Collective behaviour forms the basis for many anti-predator strategies. Within Lepidoptera, larval gregariousness has evolved convergently across many phylogenetically disparate lineages. While the selection pressures shaping variation in larval social behaviours are well investigated, much less is known about the mechanisms that control social attraction and behavioural coordination. Similarly, little is known about how secondary selection pressures associated with social living shape genome evolution. Here, using genomic data for over 60 species from an adaptive radiation of Neotropical butterflies, the Heliconiini, in which gregarious behaviour has evolved repeatedly, we explore the molecular basis of repeated convergent shifts towards gregarious larvae. We focus on three main areas of genomic evolution: differential selection on homologous genes, accelerated rates of evolution on non-coding regions of key genes, and differential gene expression in the brains of solitary and gregarious larvae. We identify strong signatures of convergent molecular evolution, on both coding and non-coding loci, in Heliconiini lineages which evolved gregarious behaviour. Molecular convergence is also detected at the transcriptomic level in larval brains, suggesting convergent shifts in gene regulation in neural tissue. Among loci showing strong signals of convergent evolution in gregarious lineages, we identify several strong candidates linked to neural activity, feeding behaviour, and immune pathways. Our results suggest sociality profoundly changes the selection pressures acting on multiple physiological, immunological and behavioural traits.

## Introduction

The convergent evolution of similar phenotypes has been a focus of biological research for over a century. Modern sequencing tools have recently allowed the exploration of the genetic bases behind convergent evolution, greatly improving our understanding of the phylogenetic relationships between complex traits such as behaviours^1,2^. Molecular changes associated with the evolution of one such behaviour, sociality, have received a great deal of attention. Across taxa, individuals living as part of a group are likely to face similar trade-offs, which potentially involve similar molecular functions^2,3–5^. These may include both a molecular response to group living, and the molecular control of the behaviour itself.

Social behaviour can impact a range of selection regimes, which likely shape patterns of genetic divergence and gene expression between solitary and social species. Aggregating introduces stresses like crowding and an increased risk of starvation and pathogen infection^6–10^, driving changes in gene expression to mitigate these costs^10,11^. Group-living animals may be driven to develop more elaborate neural circuitry to process frequent, and potentially complex, social encounters and sensory information in the form of communication signals from group members^12^. Some molecular functions may also be inherently better suited to producing the behavioural phenotype^2–4^, such as those which can be altered without negative pleiotropic effects, or only require slight modifications to produce the new phenotype^3,4^. As such, convergent behavioural evolution may involve similar mechanisms, particularly among close relatives, as the sensory cues available, sensory systems and downstream neural pathways may have similar biases and variation, and these molecular pathways may require relatively fewer modifications to produce a complex trait such as social behaviour.

To examine the extent that convergent behavioural changes involve similar molecular mechanisms we focus research on larval gregariousness in Lepidoptera. These larvae vary widely in the size, longevity and nature of their social groups, and exhibit trail-following and synchronised defensive actions that require group coordination^13–16^. Allied to these behavioural traits is huge diversity in morphology, hostplant ecology, life history and development. This diversity positions larval Lepidoptera as an informative window into the convergent evolution of group living. Gregariousness behaviour in larvae has evolved convergently across many phylogenetically disparate lineages of Lepidoptera^17–19^. Yet, to date, research into the evolution of social behaviour in larval Lepidoptera has focused almost completely on ultimate causes and ecological contexts^20–23^, with little consideration for the genetics driving the behaviour. This previous work highlights the phenotypic and environmental conditions that are important for gregariousness to evolve^19,23,24^, yet almost nothing is known of the genomic alterations required to produce the behaviour, or the effect that the behaviour-induced selection regimes have on the genome.

Here, we focus on the Heliconiini, a diverse tribe of Neotropical butterflies across which larval gregariousness has evolved independently on at least seven separate occasions^24,25^, as a potentially powerful system in which to investigate the molecular basis of social behaviour. Using genomic data for 53 species in the Heliconiini tribe and additional closely related taxa^26^, we conduct the first phylogeny-wide study testing whether convergence in social behaviours is underpinned by a molecular convergence and shared selection regimes. We examine shifts in selection pressure on orthologous genes across Heliconiini lineages with gregarious and solitary larvae, as well as testing for accelerated rates of evolution in non-coding regions associated with key loci. Finally, we compare gene expression profiles of three Heliconiini species with gregarious larvae, from three separate clades, in relation to three related species with solitary larvae, creating three monophyletic pairs of social and solitary species.

## Methods

### i) Detecting signatures of selection and convergence associated with the gregarious phenotype

To detect a signal of association between the gregarious phenotype with a shift in selective pressure (d*N*/d*S* or *ω*) across protein-coding genes (PCGs), we used the 3393 single-copy orthologous groups (scOG) list across the Heliconiinae phylogeny from Cicconardi *et al*.^26^, and a set of computational approaches designed to explore the selective landscape operating across the genome.

The first method, Branch-site Unrestricted Statistical Test for Episodic Diversification for PHenotype (BUSTED-PH), is a method to test for evidence of episodic diversifying selection associated with a specific feature/phenotype/trait^27,28^. The method first tests whether both background branches and test branches experience episodic positive selection, then checks if there is a difference between the two partitioned *ω*. We specifically tested whether there is a difference between lineages associated with the gregarious phenotype *vs* all solitary lineages. The resulting *p* values were corrected for multiple testing with the Benjamini-Hochberg method (“BH” aka “FDR”) as implemented in the R package P.ADJUST {STATS}, using an alpha threshold of 0.05. Because the within group test of episodic selection gives more stratified results rather than just differentially selected or not, we divided scOGs into two groups, *associated with gregariousness* if: there is a difference of selection between gregarious and non-gregarious (Adjusted *p* < 0.05), but not if episodic positive selection occurred in both gregarious and non-gregarious (*p* < 0.01); and *not associated with gregariousness* for the other cases. In other words, we exclude genes under selection in both gregarious and non-gregarious lineages. The two groups were plotted comparing their reciprocal gregarious *ω* vs. solitary *ω*, and their relative scaling coefficients and intercepts were analysed with SMATR^29^.

We also tested whether scOGs experienced relaxation or intensification by comparing gregarious vs solitary lineages with RELAX^30^, as implemented in the HYPHY batch language^28^. More specifically, RELAX tests whether selection pushes all *ω* categories away from neutrality, *i.e.* intensification (1 > *ω* > 1), or towards neutrality, *i.e.* relaxation (*ω* = 1). A *k* parameter is computed, which is lower than one (*k* < 1) in cases where the model supports the relaxed scenario or higher than one (*k* > 1) where intensification is better supported. Also here, *p* values were corrected for multiple testing with the Benjamini-Hochberg method, and *p* adjusted < 0.05 were considered as significant.

Finally, we used CSUBST^31^ to derive observed (*O_C_^N^* and *O_C_^S^*) and expected (*E_C_^N^* and *E_C_^S^*) numbers of non-synonymous and synonymous convergence and evaluate their rates (*dN_C_* and *dS_C_*), *ω_C_* metric, in branch combinations in a phylogenetic tree. We ran the analysis twice, once accepting all convergent events that occurred only in gregarious lineages with a threshold of *ω_C_* > 1.0 and *dN_C_* > 1.0, as an overall signal of convergence, and again with a more stringent threshold of *ω_C_* > 5.0 and *dN_C_* > 2.0 for an *arity* ≥ 2 (degree of combinatorial substitutions or the number of branches to be compared) to be considered convergence. Other statistical tests such as the one-sided Wilcoxon rank-sum tests were used to test for differences in the distributions, and the Fisher-exact tests to test whether the overlap between gene sets deviates from a random distribution. Both statistical tests were run as implemented in R and SCIPY.STATS.

### ii) Annotating conserved non-exonic elements

To identify conserved non-exonic elements (CNEEs), we used the 63-way whole genome alignment generated by Cicconardi *et al*.^26^, together with the model of neutral evolution computed with PHAST V1.4 package^32,33^. Because the majority of gregarious lineages belong to the genus *Heliconius,* we recomputed the conserved and accelerated models using PHASTCONS with the *H. melpomene* genome as reference; combining the conserved and non-conserved models with PHYLOBOOTS and using the averaged models to predict conserved elements in PHASTCONS, using default parameters. After the initial estimation of conserved elements, we merged regions within 5 bp of each other into a single conserved element. We then excluded regions: *i*) shorter than 50 bp; or *ii*) with data for less than 50 species for that region; and *iii*) with 50% of gaps in the consensus sequence; following the protocol used in Cicconardi *et al*.^26^. We used PHYLOACC-GT^34^, which computes the maximum a posteriori (MAP) Z matrix (matrix of latent states), and two Bayes factors to test for acceleration at the *Heliconius* stem. A Bayes Factor 1 (BF1), defined as the Bayes factor that compares a null model (no acceleration allowed on any branch) to the test branch model (acceleration allowed only on *Heliconius* branches) (M1); and a Bayes Factor 2 (BF2) defined as the Bayes factor comparing the test model to the full model (acceleration allowed on any branch) (M2). From all accelerated CNEEs (aCNEEs) under the M1 model we defined as *strict* those which affect three different clades of gregarious species (*e.g.*: Doris, Sara/Sapho and *Dione julia*). We also used PHYLOP from PHAST V1.4 package^32,33^ [--mode CONACC --wig-scores --method LRT] to generate the acceleration/conservation scores.

### iii) Enrichment among CNEEs

To test for gene ontology (GO) terms of functional elements enriched in Gregarious-accelerated CNEEs, we used the approaches as implemented in Cicconardi *et al*.^26^, *i*) two permutation approaches to account for possible biases towards particular gene functions; *ii*) a genomic fraction approach as implemented in GREAT^35^. To each gene in the reference genome annotation (*H. melpomene*) was assigned a regulatory domain as the 5 + 1 kb with extension strategy, similarly to GREAT^35^. In brief, to each coding locus a regulatory domain is assigned consisting of a basal domain that extends 5 kb upstream and 1 kb downstream from its transcription starting site (TSS), which may overlap between other basal domains of flanking loci, plus a further extension up to the basal domain of the nearest upstream and downstream locus, up to 1 Mb.

The “simple permutation test” was computed calculating the expected probability of overlap between the regulatory domains of genes of a specific GO term and 10,000 random aCNEEs datasets, shuffling the CNEEs and randomly selecting the same amount of accelerated CNEEs among all CNEEs, using a binomial test to generate a *p* value. For the second test “Genomic region test”, similarly to GREAT, the binomial test was executed over the total fraction of genomic regions associated with a given GO term. A third method was implemented “Reshuffling permutation set” in which BEDTOOLS SHUFFLE [-excl gff3 -chrom - chromFirst-noOverlapping] was adopted to shuffle all detected aCNEEs across the entire genome avoiding overlaps with annotated genes and comparing the overlap with putative regulatory elements, in a binomial test framework. Multiple test correction was done with the Benjamini–Hochberg FDR (FDR_bh) and Bonferroni as implemented in python library STATSMODELS.STATS.MULTITEST. With both approaches we tested all biological processes where at least two aCNEEs were overlapping with a regulatory domain of the given GO term. We used REVIGO^36^ to summarize results.

In the same fashion we also looked for genomic regions, gene annotations and gene sets (BUSTED-PH) enriched for aCNEEs. To do this, we generated non-overlapping 100 kb sliding windows and computed the probability of observing aCNEEs based on the binomial distribution over the 10,000 permutations of randomly selected accelerated CNEEs among all CNEEs, where the number of trials is the number of aCNEEs in the window and the probability of success is the average of the permutations; the excess of aCNEEs per regulatory domain using the distribution over the 10,000 permutations as implemented in Cicconardi *et al.*^26^; and the excess of aCNEEs to the gene of differentially selected scOGs (BUSTED-PH) over 10,000 permutations considering as background all the genomic regions/regulatory domains of all the tested scOGs. The *p* values of those tests where then corrected as before.

### iv) Larval rearing and sample collection

Six species from the Heliconiini tribe were used: *Dione juno, Heliconius doris* and *H. sara*, which are all gregarious as larvae, and *Agraulis vanillae, H. hecale* and *H. erato*, which are solitary. These six species belong to three distinct monophyletic pairs of gregarious and solitary species and were used to contrast gregarious lineages to solitary close relative species. Although behavioural variation exists among gregarious species, we selected these species for comparison because all three remain gregarious up to, and often including, pupation^37–39^. Stock populations of each species were established from wild caught females collected around Gamboa, Panama, in 2013. All larvae are therefore first or second generation, insectary reared individuals, with the exception of the locally less abundant *D. juno*, which was opportunistically sampled from a single brood found as eggs on a *Passiflora vitifolia* in the outdoor host plant stocks. Stock populations were held in ambient, tropical conditions outside in ∼ 2 × 2 × 3 m cages, were fed on artificial feeders, and for *Heliconius spp* with ∼20% sugar/bee pollen solution with supplemented floral resources (*Lantana sp.* and *Psiguria sp.*). Each species had access to their preferred host plant (*D. juno* and *H. hecale*: *Passiflora vitifolia*; *H. doris*: *P. ambigua*; *H. sara*: *P. auriculata*; *A. vanillae* and *H. erato*: *P. biflora*). Eggs were collected daily, and larvae were subsequently reared on shoots in pop-up cages at ambient conditions. All species were reared concurrently so were exposed to common environmental conditions. Larvae were sampled in the early 5th instar, before the prepupal instar, or “wandering stage”, where behaviour often changes drastically as larvae cease foraging and search for a pupation site (*e.g.*: Smallegange *et al*.)^40^. We used five separate individual samples per species, giving a total of 30 samples. For the gregarious species, *H. doris* and *H. sara* samples were taken from individuals from four separate broods, whereas all five *D. juno* samples were taken from the same brood. Solitary species were selected from the same generation, but as eggs were derived from stock cages with multiple fertilised females, the kinship between individuals was not known. The brain and connecting suboesophageal ganglion were dissected out in RNA*later* (which stabilises RNA, preventing its breakdown; ThermoFisher Scientific), and stored at -20°C.

### v) RNA extraction and sequencing

We extracted total RNA from each sample consisting of an intact, attached brain, and suboesophageal ganglion following the instructions of the RNeasy micro kit according to the manufacturer’s protocol (Qiagen). RNA concentration, purity and integrity were assessed on an Agilent 4150 TapeStation (Bristol Genomics Facility, UK). To improve 28S RNA peak visibility, samples were not subjected to heat denaturation treatment, following the suggested approach for assessing insect RNA^41^. All samples had an RNA Integrity Number (RIN) of >= 9, indicating that the RNA was of suitable quality for sequencing^42^. Samples were sent to Novogene (UK) for library preparation (with poly A enrichment) and transcriptome sequencing. Libraries were sequenced on an Illumina NovaSeq 6000 platform, producing approximately 150 bp paired-end reads. We quality checked raw reads using FASTQC v. 0.11.9^43^ and trimmed them using TRIMMOMATIC v. 0.39^44^ to remove adapter contamination, low quality reads, and reads shorter than 75 bp, using default command line settings.

### vi) Differential gene expression analysis

Annotated genomes for all six Heliconiini species were obtained from Cicconardi *et al.*^26^. We used STAR v. 2.7.11a^45^ to map trimmed reads to genomes and HTSEQ v. 2.0.3^46^ to count the mapped reads per gene. A table of Heliconiini orthologous groups (OGs) was obtained from Cicconardi *et al.*^26^ and filtered to retain only scOGs containing genes for all of our target species (total retained genes = 6885). We then matched the read counts of genes to their corresponding OGs in order to compare OG expression across the study species.

All analyses of differential gene expression were performed in R v. 4.1.2^47^ using LIMMA v. 3.5.3^48^. Raw read counts were first normalised based on gene length, library size and GC content using CQN v. 1.4.0^49^, to account for potential bias due to differences in homologous gene lengths between species^50^.

We compared gene expression between species in three separate pairs based on larval social behaviour and clade, where each pair consisted of a gregarious and solitary species belonging to the same clade: i) *D. juno* vs *A. vanillae* (diverged ∼14 mya; ii) *H. doris* vs *H. hecale* (diverged ∼9 mya), and iii) *H. sara* vs *H. erato* (diverged ∼5 mya)^26^. Additionally, we performed a full comparison of all three solitary species *vs* all three gregarious species. Differentially expressed genes (DEGs) were selected using adjusted *p* value threshold of 0.05. Of these, we selected only those with a minimum log_2_ fold change (log_2_FC) of ±1 to account for any differences in brain tissue scaling (overall size, region volumes etc.) between species^51^.

### vii) Functional enrichment and overlap of DEGs

A list of GO terms corresponding to our OGs was obtained from Cicconardi *et al.*^26^, providing information on the biological function towards which each DEG is likely to contribute^52^. These GO terms were used for functional enrichment analyses, using the R package topGO v. 2.46.0^53^ on each species pair, with the aim of highlighting the functions that are overrepresented in our gene lists of interest, both down and up regulated.

Finally, we examined the overlap of DEGs between the three species-pair comparisons to identify convergent shifts in expression. We used 10,000 iteration permutation tests to assess the probability that the number of shared DEGs between the two- and three-way species-pair comparisons is greater than expected by chance. In detail, n genes were sampled without replacement from the full list of OGs, where *n* = the number of observed DEGs from a given species pair, to create a list of random DEGs. The number of shared genes between both (or all three) randomised lists was recorded, and the process repeated over 10,000 iterations to give a distribution of 10,000 sets of shared genes. We then tested the observed count of shared DEGs against this distribution for the probability of it occurring by chance.

## Results and Discussion

### i) Convergent shifts in selection regimes shapes coding genes in gregarious species

Using a set of single-copy orthologous groups (scOGs) obtained from 53 Heliconiinae species (Figure 1)^26^, we first explored the strength of convergent evolution among protein coding genes (PCGs) between gregarious species. To do so, we adopted three analytical approaches: *i*) using BUSTED-PH^27,28^ we tested for evidence of different selective regimes between solitary and gregarious lineages; *ii*) using RELAX^30^ we tested whether these shifts in selection regime can be explained with a relaxed selection pressure (a shift of rate-classes towards neutrality; *k* < 1), or with an intensification of selection pressure (a shift of rate-classes towards purifying and positive selection; *k* > 1); finally, *iii*) we scanned scOGs for signs of convergent substitution with CSUBST, a method for detecting the Combinatorial SUBSTitutions of codon sequences^31^. The CSUBST algorithm scans all combinations of target lineages from pairwise (*arity* = 2) to multiple (*arity* > 2) comparisons to identify genes that are under convergent positive selection (*ω_C_* > 1.0).

**Figure 1.**
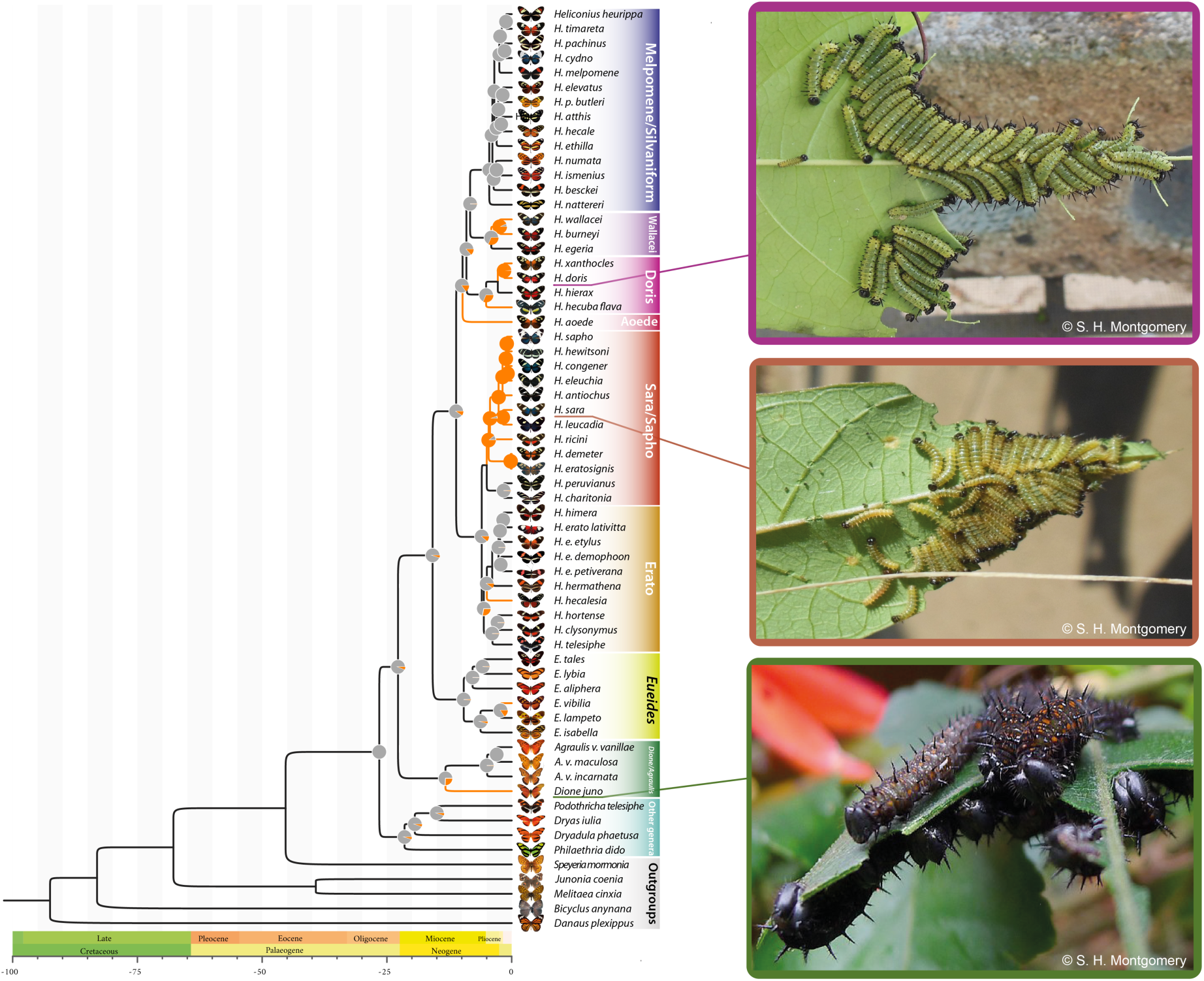
Heliconiini phylogeny and ancestral state reconstruction of gregarious behaviour. (A) Dated phylogeny of Heliconiini species from Cicconardi *et al.*^26^. Orange branches indicate lineages having gregarious behaviour, while pie charts indicate the maximum likelihood estimate of the most recent common ancestor to have gregarious larvae from McLellan and Montgomery^24^. (B) Photographs illustrating independently evolved larval social groups in (from top to bottom) *Heliconius doris*, *Heliconius sara*, and *Dione juno*.

Our BUSTED-PH^27,28^ analysis detected 203 scOGs (6%) to be under distinct selection regimes in gregarious lineages (adjusted p values < 0.05; Figure 2A,F; Table S1). While the median strength of selection acting on scOGs in solitary (*ω_s_*) and gregarious lineages (*ω_g_*) is only marginally different overall (0.072 for not differentially selected genes (nDSGs) vs 0.066 for DSGs; One-sided Wilcoxon rank-sum test p adj = 0.047), we detect a much higher median *ω_g_* for those genes with distinct selection regimes between solitary and gregarious lineages (0.12 vs 0.16; One-sided Wilcoxon rank-sum test p adj 2.2×10^−16^; Figure 2A). This is associated with a proportion of nucleotides evolving under strong purifying selection in the solitary lineages, which shift towards neutrality or positive selection in gregarious lineages (Figure 2A, insert). Our RELAX analysis supports a role for increased neutrality in a class of scOGs in gregarious lineages, with 36% (1224 scOGs) showing a signature or relaxation (Table S1). Among this class of genes, no gene ontology (GO) terms are significantly enriched but several are associated with the response to viruses, including a *Lachesin*-homolog gene and other ubiquitine-related genes (*e.g. sina*, an E3 ubiquitin-protein ligase). In other insects, viruses have been demonstrated to interact with *lachesin* proteins during their attachment to neurons^54^, and ubiquitination is known to play an important role in viral infection^55^. In *Bombyx mori* a *sina*-homolog is downregulated to inhibit cellular interactions with viral proteins (*e.g. BmNPV*)^56^. A smaller proportion of scOGs also show evidence of intensified positive selection in gregarious lineages (∼4%, 122 scOGs; p adj < 0.01; Figure 2C, F). Among these genes, an odorant receptor (OR43; ortholog of the *B. mori* OR49^57^) is known to bind to the antimicrobial and larval food attractant compound citral^58,59^.

Our CSUBST^31^ analysis next sought to identify convergent substitutions in gregarious lineages. We first assessed the degree of convergent selection in genes previously identified as evolving under differential selection (BUSTED-PH), relaxation or intensification (RELAX). To do so, we selected a relatively low threshold for convergence of selection (*ω_C_* > 1.0) and rate of non-synonymous convergence (*dN_C_* > 1.0). Genes under differential selection showed a strong increase in both *ω_C_* and the rate of non-synonymous convergent substitutions (medians: 325.9 *ω_C_* and 11.6 *dN_C_*) compared with other scOGs, where no differential selection was detected (Medians: 26.3 *ω_C_* and 7.7 *dN_C_*; One-sided Wilcoxon rank-sum test p adj value < 2.7×10^−5^), with no significant difference in the rate of synonymous convergent substitutions (*dS_C_*), which can function as negative control (Figure 2B). For scOGs under relaxed and intensified selection there is less enrichment of for signals of convergence, but in gregarious lineages, scOGs under intensified positive selection show a stronger enrichment of *ω_C_* compared with scOGs under relaxed selection (*ω_C_* medians: 57.9 *vs* 22.8; One-sided Wilcoxon rank-sum test p adj < 3.0×10^−3^). This pattern suggests that the intensified positive selection in gregarious lineages is partly driven by convergent substitutions. Finally, selecting a more stringent threshold of *ω_C_* > 5.0 and *dN_C_* > 2.0 for an *arity* ≥ 2, we were able to identify 192 scOGs under convergent positive selection (*ω_C_*^+^) (Figure 2E, F). Most of this convergence occurred between pairs of gregarious lineages (*arity* = 2) (Figure 2E). The most frequent lineages under *ω_C_*^+^ were *Heliconius aoede* and *Dione juno* (32 scOGs with median *ω_C_* of 9.9), which split from their MRCA more than 25 mya (Figure 1). Among these genes we find *Herc4*, a ubiquitin ligase4, *Notch,* and *Neurotrophin 1* (*NT1*), which are all connected with Toll-related receptors involved in host defense^60^, and a (cytosine-5) tRNA methyltransferase (*Mt2*), which in *Drosophila* is implicated in the innate immune response and lipid homeostasis^61^. *Heliconius aoede* also has the highest number of scOGs under *ω_C_*^+^, with over 100 genes, followed by *D. juno* and *H. ricini* (Figure 2e). The scOGs implicated in convergence among the highest number of lineages are a *Rcc1*-homolog and *kazachoc* (*kcc*) (Figure 2E). *Rcc1* has roles in cell proliferation, cell survival, apoptosis, epigenetic regulation, and neuronal specification^62,63^. *Kazachoc* encodes a potassium:chloride symporter expressed in the insect central nervous system and is implicated in modulating neural firing rates^64^.

Exploring overlap between genes highlighted by the independent analyses above, we find significantly more overlap than expected by chance in four of the six pairwise datasets (100,000 permutations; p values < 0.008). The most interesting overlap between genes under relaxation, differentially selected, and under *ω_C_*^+^ include 14 scOGs. Among them are *CycB3*, *Glg1*, *Vps51*, *Fancl*, which are implicated in the regulation of cell cycle, Golgi apparatus, and response to DNA damage stimuli^65–68^, and *norpA*, which in *Drosophila* plays a prominent role in the transduction of light and environmental stimuli^69^. Overall, our analysis of protein coding genes suggests a shift towards gregariousness that affects the selection regimes shaping multiple biological processes, with a strong signal of convergence among a small subset of genes.

**Figure 2.**
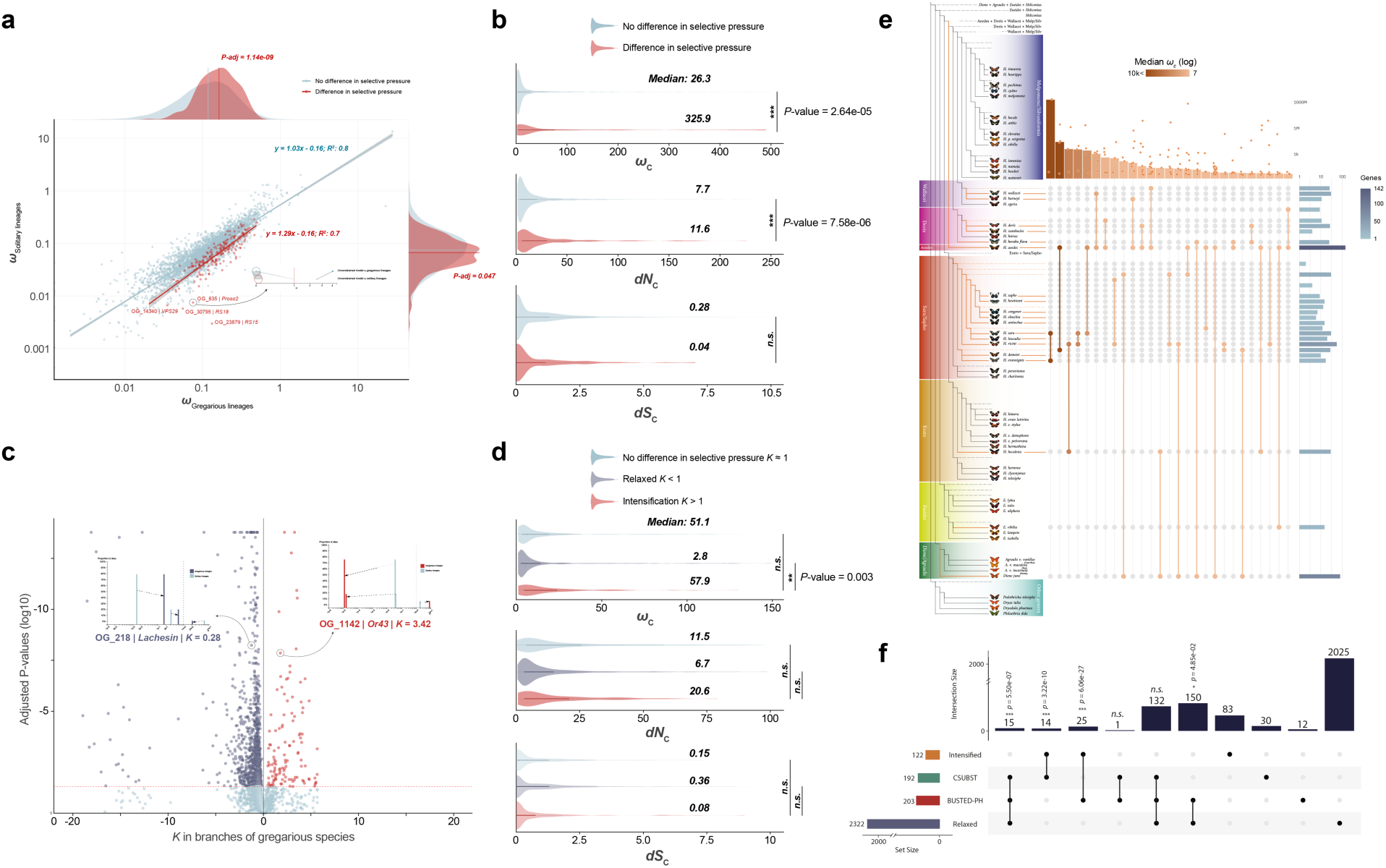
Convergence of molecular evolution in single-copy orthologous groups and their association with gregarious lineages. (A) Scatter plot of scOGs showing differential selecting regimes between gregarious and solitary lineages. Red dots correspond to genes that are differentially selected in gregarious lineages compared with solitary lineages, while light blue dots are not. Regression lines show significant slops. Along *x* and *y* axes the distributions of *ω* in gregarious and solitary lineages for the genes associated with gregarious and solitary lineages are plotted. The insert shows the three-rate classes of *ω* between gregarious vs solitary lineages in *Prosα2*, as an example. (B) Violin plots of convergent positive selection (*ωC*), rate of non-synonymous convergent substitutions (*dNC*), and rate of synonymous convergent substitutions (*dSC*) between differentially (red) and not-differentially selected genes (light-blue). (C) Scatter plot of RELAX analysis. Light-blue dots represent genes under relaxation (*k* < 1; p adj. < 0.05); while red dots correspond to genes under intensification (*k* > 1; *p*-adj. < 0.05). *x*-axis is log2 transformed to centre *k* = 1 to 0. The red-dashed line corresponds to the *p* adj threshold of 0.05. The two inserts show the change in the three-rate classes of *ω* between gregarious vs solitary lineages in *Lachesin* and *Or43* loci, which are relaxed or intensified, respectively, as an example. *Lachesin* has been associated with change in crawling behaviour in *Drosophila*, while *Or43* is an odorant receptor known to be able to bind to the antimicrobial and larval food attractant compound citral. (D) Violin plots of convergent positive selection (*ωC*), rate of non-synonymous convergent substitutions (*dNC*), and rate of synonymous convergent substitutions (*dSC*) between relaxed (light-purple), intensified (red), and no-difference genes. (E) CSUBST analysis across Heliconiini phylogeny. Next, the expanded tree diagram shows the lineages involved in convergent positive selection (*ω^+^C*). The histogram on top shows the median value of *ω^+^C* for specific pairs. On the right-most side the histogram shows the number of genes evolving convergently in the specific branch. (F) UpSet plot showing the intersection of the four gene lists. On top of the vertical histogram the corresponding p values of the probability of the overlap being due to a random distribution. In all plots asterisks the size of the p value: *n.s.* = not significant; * < 0.05; ** < 0.01; *** < 0.001.

### ii) A subset of conserved non-exonic elements show accelerated rates of evolution in gregarious lineages

To determine the extent of convergent molecular evolution at non-coding loci, we next compiled a total of 148k conserved non-exonic elements (CNEEs) across the 63-way genome alignment^26^, using *H. melpomene* as a reference. CNEEs are small genomic regions previously shown to be enriched with *cis*-regulatory elements^26,70^. Within these 148k CNEEs, we explored convergent shifts in molecular rate associated with transitions to gregarious behaviour^71^ (Figure 3). We identified 15.3% of CNEES that show variable rates of evolution across the phylogeny (acceleration under a “full model” M2 has a higher probability, expressed as Bayes Factor cutoff 1.0, than “null model” M0) and 9.8% that show convergent acceleration (aCNEEs) in gregarious lineages (acceleration under a “target model” M1 has a higher probability than M0 and M2), corresponding to ∼10% (1.32Mbp, 14.6k CNEEs) of the reference genome. Of these regions roughly a third (29.2%) of all aCNEEs are convergently accelerated in at least three different clades of the phylogeny (Figure 3A, B).

We found 37 candidate genes associated with gregarious behaviour that harbour more aCNEEs than expected by chance (Figure 3A, D; Table S3; p adj. < 0.05). These include the putative *phenylalanyl-tRNA synthetase β-subunit homolog* (*β*-PheRS). *β*-PheRS is an important enzyme that acts by charging tRNAs with their cognate amino acid and, in *Drosophila*, manipulating *β*-PheRS expression affects growth speed, the timing of pupation, and behaviours including feeding and roaming^72^. More broadly, aCNEEs are enriched in putative regulatory domains of genes associated with more than 40 GO terms (Figure 3C; Table S4). The most significantly enriched GO terms include those associated with viral detection, defence response to oomycetes and fungi, regulation of antimicrobial peptide production, and Toll signalling pathway - the innate immune response pathway in insects.

Finally, we explored the intersection between CNEEs and scOGs and found that in putative regulatory domains of the scOGs, 15% of the CNEEs are accelerated (M2), and 9.4% are convergently accelerated (M1), of which 27% are accelerated in at least three clades (Figure 3E). These are similar numbers to our genome-wide analyses, suggesting that our scOG dataset is a robust representation of the entire coding gene repertoire. Focusing on scOGs with putative associations with gregarious behaviour (*i*.*e.:* differentially selected, relaxed, intensified and positively converging genes), we found a significant enrichment of strict aCNEEs (*p* value < 0.05; 34.6% of CNEEs) in the putative regulatory domains of scOGs with differential selective regimes in gregarious lineages. This translates to a two-fold density of strict aCNEEs in the differentially selected scOGs (0.4 aCNEE/scOG) compared with the background scOGs (0.2 aCNEE/scOG). We also observed that aCNEEs in putative regulatory domains of differentially selected scOGs had higher acceleration rates (median: 1.31 vs 1.28) and a lower conservation rate (median: 0.091 vs 0.093) (Figure 3F). Overall, our analyses of non-coding genomic regions show a strong signal of convergence associated with gregarious larvae. Importantly, the combination of analytical approaches applied to both coding and non-coding regions allow us to detect that selection for gregarious behaviour is simultaneously acting at the same loci: in protein coding regions, by changing amino-acid composition (purifying and/or positive selection), and through accelerated rates of sequence evolution in *cis*-regulatory elements (non-coding elements).

**Figure 3.**
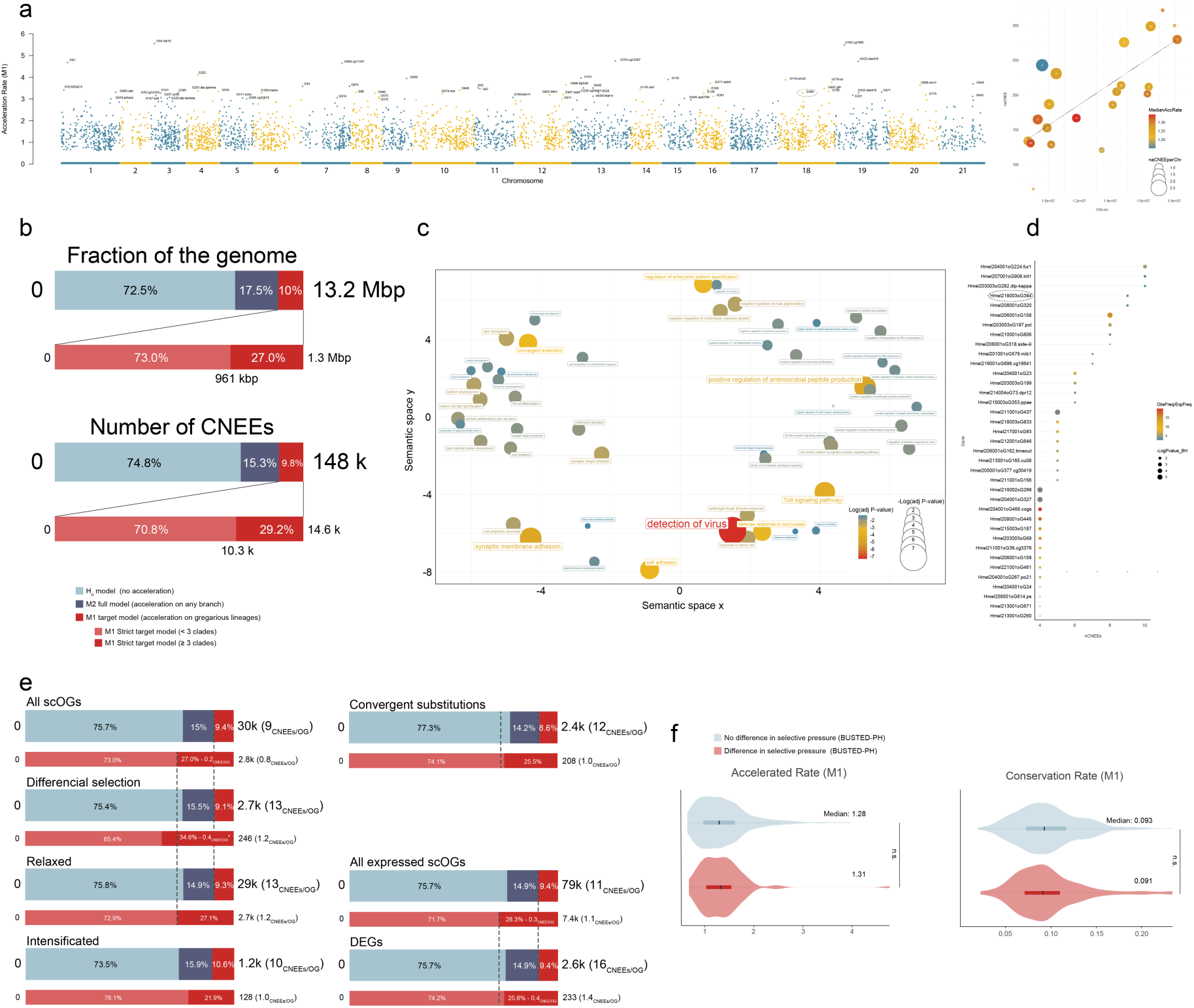
Convergent evolution of conserved non-coding elements in gregarious lineages throughout Heliconiini phylogeny. (A) Distribution of “strict” aCNEEs across the genome of *H. melpomene*, used as reference. *y*-axis is the acceleration rate under the M1 model. On the left side the positive correlation between the number of aCNEEs and chromosome lengths. Red-shift shows an enrichment of high rate. In particular chromosomes 11 and 14 show on average higher rates, while chromosome 4 show an excess of aCNEEs with respect its size, but with low rate on average; while chromosome 21, the sexual chromosome (Z), and 2 show a very low number of aCNEEs. (B) Staked histograms showing the proportion of the CNEEs as fraction of the genome (up) and in terms of model distribution (down). (C) Semantic plot of the enriched GO terms using genes interested by *strict* aCNEEs present in putative regulatory elements (see Methods). Large and red-shift circles correspond to GO terms with have smaller *p* values. (D) Bubble plot of some of the 73 genes enriched with aCNEEs in their regulatory elements (Table S3). Circle size shows the log10 of the *p* adj. values of the enrichment, while the red shift shows higher values of the ratio between the observed vs the expected frequency. Genes that are also differentially expressed in at least one comparison are listed in bold. (E) Staked histograms showing the proportion of the different classes CNEEs in different gene sets, including expressed and differentially expressed scOGs (DEGs). (F) Violin plots of accelerated rate and conservation rate for differentially selected genes (BUSTED-PH gene set). In all plots asterisks the size of the *p* value: *n.s.* = not significant; * < 0.05.

### iii) Shifts in gene expression during the independent evolution of gregarious behaviour

To further explore the impact of convergent shifts towards gregarious behaviour on neural gene expression, we collected larval brain transcriptome data for three phylogenetically independent pairs of solitary/gregarious species (Figure 4A), with five replicates per species. Data normalisation, aligning gene lists between species pairs, and removing genes which had zero reads across all species resulted in a final dataset of 4,874 scOGs shared across all six species (obtained from Cicconardi *et al.*^26^). We next compared patterns of gene expression between each pair of solitary/gregarious species. This revealed a relatively high number of scOGs with significant differential expression within each pair (Figure 4B-D; Table S5), with qualitatively greater divergence with increasing phylogenetic distance (Figure 4A), suggesting a role of genetic drift in shaping gene expression divergence. Separate tests for functional enrichment on differentially expressed genes (DEGs) between species pairs revealed no significantly enriched GO terms after Benjamini-Hochberg p value correction for false detection rate (all *p* > 0.05, Table S6).

We next examined the amount of DEG overlap between pairs. Permutation tests revealed that the level of DEG (up- and downregulated in gregarious lineages) overlap between *D. juno* and *H. doris* is significantly greater than expected by chance (47% and 56% to total genes, respectively; *p* < 0.001, Figure 4E, Table S7), as is the degree of overlap at the intersect between all three pairwise comparisons (23%, 28%, and 26% in *D. juno, H. doris* and *H. sara*, respectively; *p* < 0.001, Figure 4E, Table S7). The higher strength of overlap between *D. juno* and *H. doris* is notable, as the larvae of these species are more strictly gregarious throughout larval development, relative to *H. sara*^37,38^. Considering overlapping up- and downregulated genes separately, the level of upregulated gene overlap between *D. juno* and *H. doris* is again significantly greater than expected by chance (29% and 30%, respectively; *p* = 0.002, Table S7), and the same is true for the upregulated genes at the intersect between all three pairs (10%, 10%, and 9% in *D. juno, H. doris* and *H. sara*, respectively; *p* = 0.008). In contrast, overlap among downregulated genes does not deviate from random expectation (Table S7). Considering scOGs with at least three aCNEEs in their putative regulatory regions, we identified 14 of them that are differentially expressed between one or more pairs of solitary and gregarious Heliconiini (Table S3). Six of these harbour more aCNEEs than expected by chance (Table S3; *p*-adj. < 0.05). Of particular note is *dpr12* (Figure 3D), which is differentially expressed in all three species pairs, and is involved in dopamine signalling, a neuromodulator with known regulatory roles in several insect behaviours linked to sociality, including feeding behaviour and aggression (reviewed in Verlinden^73^); and two *semaphorins* (*sema-1a and sema-1b*, the former is shown in Figure 4F), which is involved in growth cone guidance through its role in axonal repulsion^74^.

Several of the convergent DEGs with the greatest mean log_2_ fold-change identified in our study are predicted to be involved in functions putatively linked to social behaviour, such as hormonal pathways (*5htr*, *dop2r*), sensory system development and function (*CG31559, ap-1σ, hdc, dan*), immune response (*rnf19b, CG3829, samdc, senju, ubc7*), feeding behaviour and metabolism (*CG3552, mthl3, rmi1, hcrtr2, CG30344, indy*), and neural development (*tao*) (Table S8). While genes associated with sensory functions may provide clues as to the stimuli detected by social larvae, the convergent upregulation of genes controlling hormonal pathways, such as serotonin and dopamine receptors, may also suggest that modulation of how social cues are used plays an important role in the evolutionary transition to social behaviour. As observed in other species, the activity and regulation of these hormones can be crucial in the production of the social behavioural phenotype^75–77^, potentially because of their involvement in feeding^78–80^ and aggression or attraction/repulsion responses between conspecifics^75,77,81^. Elsewhere, these candidate genes may be associated with secondary selection pressures which result from social living. For example, feeding regulation is likely to be an important social phenotype given higher competition for resources between group members^7,8^ and the greater risk of starvation during collective defoliation of a host plant^82,83^. Similarly, social organisms are more susceptible to disease, given their close proximity to one another and sharing of food sources^84^, and group living is hypothesised to be associated with increased selection on individual immunity^84,85^.

**Figure 4.**
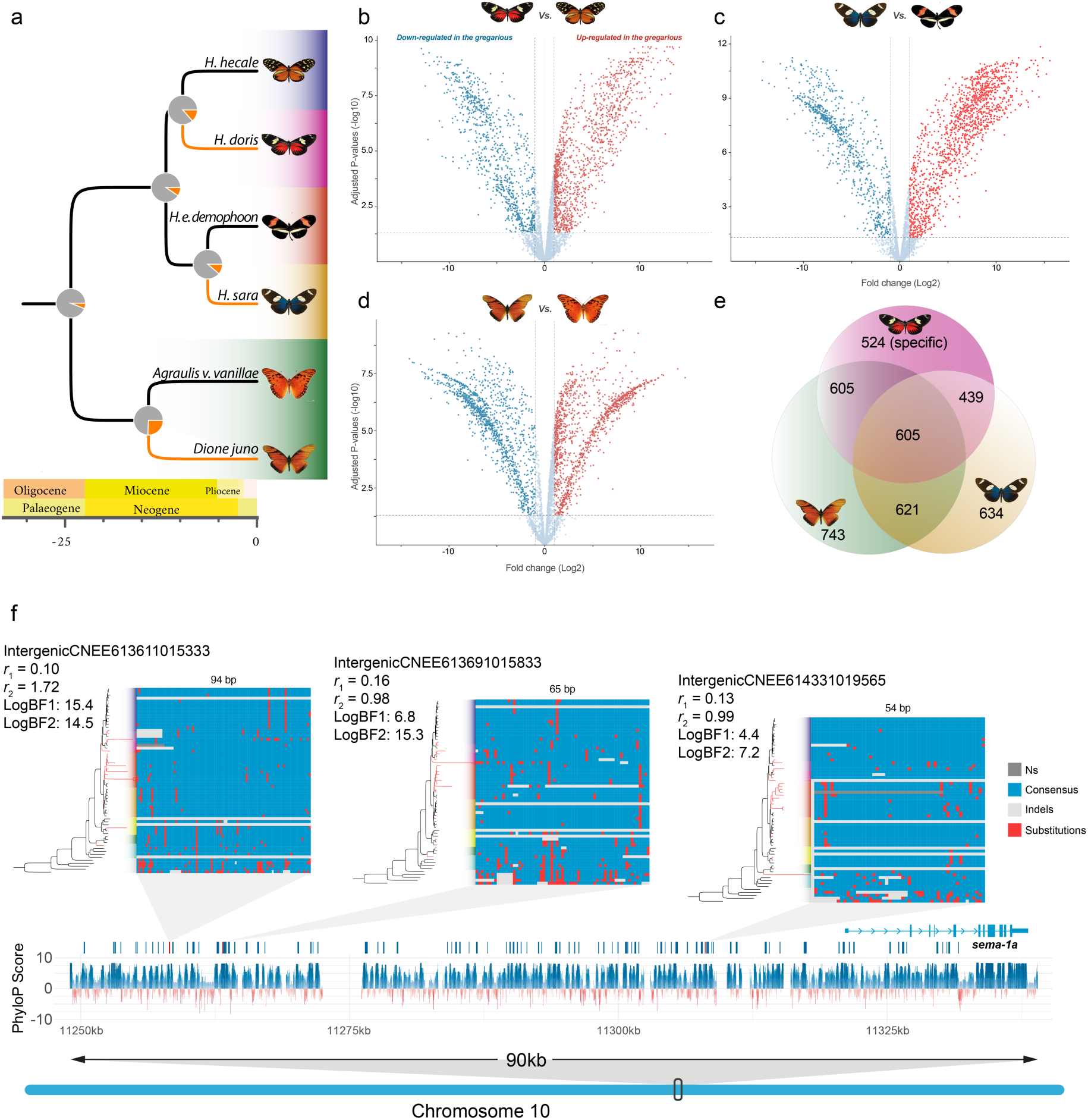
Convergent gene expression analysis across three pairs of species across the Heliconiini phylogeny. (A) Dated phylogeny of the three species pairs used as proxy to study the gene expression convergence in gregarious lineages. As in Figure 1, orange branches indicate lineages with gregarious larvae, while pie charts indicate the ancestral state reconstruction of the most recent common ancestor having gregarious larvae. (B-D) Volcano plot of the differentially expressed scOGs in the three pairs of species. In each pair the solitary larva species was used as “control”, therefore red points correspond to upregulated genes in the gregarious species, while the light-blue points represent downregulated genes. Horizontal dashed lines indicate *p* = 0.05, vertical dashed lines indicate a ±1 log2FC threshold. (E) Venn diagram of the DEGs in common in the three studied pairs. Asterisks corresponds to the size of the p value: *n.s.* = not significant; * < 0.05; ** < 0.01; *** < 0.001. (F) Genomic locus around the gene *sema-1a*. The locus has a number of conserved loci as indicated by the phastCons and PhyloP tracks. The locus has three aCNEEs for each of them a phylogenetic tree is shown with branch lengths proportional to the corresponding acceleration rates for the gregarious lineages (red branches). Next to the phylogenetic tree the alignment of the CNEE is also showed. For each aCNEE the conservation (*r1*), acceleration rates (*r2*), and their respective log of the bayes factors (logBF) are also listed.

### iv) Conclusions

Using multiple approaches across transcriptomic, protein coding and non-coding genomic data, our analyses provide a robust case that convergent transitions towards gregarious larval behaviour are associated with genome-wide convergent molecular evolution. We have identified protein-coding genes under differential selection in gregarious lineages, suggesting that changes at these loci may be important in the evolution of the gregarious phenotype. We also reveal a parallel signal of convergence at putative *cis*-regulatory elements, providing evidence of common shifts in selection affecting coding and non-coding elements of a wider locus. Finally, we have taken the first steps in exploring these gene expression profiles, revealing a convergent pattern of differential expression in association with convergence in the gregarious phenotype, with a number of genes associated with convergent aCNEEs being differentially expressed in independent gregarious lineages. Our comprehensive analyses identify the first candidate loci associated with gregarious larval behaviour in Lepidoptera, in particular highlighting the serotonergic and dopaminergic pathways as a potential mechanism for transitioning to larval gregariousness. Given the importance of these molecular mechanisms on other insect social behaviours, our data provide encouraging loci for future functional investigation. Crucially, our data also reveal the likely importance of genes involved in a wide range of physiological and behavioural traits, which likely evolved in concert with gregariousness *per se*. These include feeding behaviour, immune system function, and possibly aggression, as components of the repeated evolution of the gregarious phenotype across the Heliconiini. Little is known of the specific ‘tolerance’ genes that are likely to facilitate social behaviours^86^, and our data provide a first step towards the identification of genes involved in anti-aggression, anti-cannibalism, and the prevention of egg consumption. More broadly, our analyses highlight the rich potential of phenotypically informed comparative analyses of densely sampled genomic datasets of closely related, but phenotypically divergent, phylogenetic groups.

## ACKNOWLEDGMENTS

This work was carried out using the computational facilities of the Advanced Computing Research Centre, University of Bristol. We are grateful to Christy Waterfall at the Bristol Genomics Facility for her assistance and detailed guidance. We thank Adriana Tapia, Moises Abanto, Oscar Paneso, Cruz Batista Saez, and STRI for support at the Gamboa insectaries, Panama. This work was funded by a Royal Commission for the Great Exhibition Research Fellowship, a Short-term STRI Fellowship, British Ecological Society Research Grant (3066), and a NERC IRF (NE/N014936/1) to SHM, and an SWBio DTP Studentship to CF (BB/M009122/1).

## AUTHOR CONTRIBUTIONS

SHM conceived and designed the research, with input from FC and CFM. CFM collected and analysed data the gene expression data, with assistance from AS, following larval rearing and brain dissection performed by SHM with support from WOM. FC performed all genomic analyses. CFM and FC wrote the first draft with assistance from SHM, with subsequent edits performed equally between CFM, FC and SHM.

## Data and code availability

The code used for this study and statistical analysis are available in the Git repository: https://github.com/francicco/-ConvergentMolEvolutionGregLarvalBehaviour

